# TRPV1-mediated sonogenetic neuromodulation of motor cortex in freely moving mice

**DOI:** 10.1101/2022.10.28.514307

**Authors:** Kevin Xu, Yaoheng Yang, Zhongtao Hu, Yimei Yue, Jianmin Cui, Joseph P. Culver, Michael R. Bruchas, Hong Chen

## Abstract

**Background:** Noninvasive and cell-type-specific neuromodulation tools are critically needed for probing intact brain function. Sonogenetics for noninvasive activation of neurons engineered to express thermosensitive transient receptor potential vanilloid 1 (TRPV1) by transcranial focused ultrasound (FUS) was recently developed to address this need. However, using TRPV1-mediated sonogenetics to evoke behavior by targeting the cortex is challenged by its proximity to the skull due to high skull absorption of ultrasound and increased risks of thermal-induced tissue damage.

**Objective:** This study evaluated the feasibility and safety of TRPV1-mediated sonogenetics in targeting the motor cortex to modulate the locomotor behavior of freely moving mice.

**Methods:** Adeno-associated virus was delivered to the mouse motor cortex via intracranial injection to express TRPV1 in excitatory neurons. A wearable FUS device was installed on the mouse head after a month to control neuronal activity by activating virally expressed TRPV1 through FUS sonication at different acoustic pressures. Immunohistochemistry staining of *ex vivo* brain slices was performed to verify neuron activation and evaluate safety.

**Results:** TRPV1-mediated sonogenetic stimulation at 0.7 MPa successfully evoked rotational behavior in the direction contralateral to the stimulation site, activated cortical neurons as indicated by the upregulation of c-Fos, and did not induce significant changes in inflammatory or apoptotic markers (GFAP, Iba1, and Caspase-3). Sonogenetic stimulation of TRPV1 mice at a higher acoustic pressure, 1.1 MPa, induced significant changes in motor behavior and upregulation of c-Fos compared with FUS sonication of naïve mice at 1.1 MPa. However, signs of damage at the meninges were observed at 1.1 MPa.

**Conclusions:** TRPV1-mediated sonogenetics can achieve effective and safe neuromodulation at the cortex with carefully selected FUS parameters. These findings expand the application of this technique to include superficial brain targets.

## Introduction

The evolution of brain neuromodulation tools has provided unprecedented opportunities to probe neural circuits, understand brain function, and develop new treatment strategies for brain diseases. Transcranial neuromodulation tools, such as direct current, magnetic stimulation, and ultrasound stimulation, offer noninvasive ways to stimulate the brain and have contributed to the understanding of brain function [1,2]. The lack of cell-type specificity in these tools, however, limits their utility in understanding the brain at cellular resolution. Genetic-based neuromodulation tools, such as optogenetics and chemogenetics, encode stimulus-sensitive probes into a defined neuron population and have transformed fundamental neuroscience research [3,4]. Each method, however, suffers from its own limitations. Most commonly, optogenetics requires the invasive implantation of optical probes to deliver light to opsin-encoding neurons, limiting the ability to study the brain without the risk of ischemia and inflammation. Noninvasive optogenetics modulates the activity of opsin-encoding neurons via transcranial illumination, but light scattering in brain tissue limits its depth penetration in large animal models [5]. On the other hand, chemogenetics noninvasively activates neurons encoding designer receptors exclusively activated by designer drugs (DREADDs) via minimally-invasive systemic delivery of designer drugs, but the long residence time of circulating drugs sacrifices the temporal resolution of this technique. There is a clear need for techniques that can facilitate noninvasive, cell-type specific neuromodulation with high spatiotemporal resolution and the potential to be scaled up to large animals and humans.

Sonogenetics has great potential to fulfill this gap. Analogous to other genetic-based neuromodulation tools, sonogenetics uses focused ultrasound (FUS) to modulate the activity of neurons encoding ultrasound-sensitive actuators [6]. Unlike other stimulation modalities (e.g., light, electricity, and magnetic fields), FUS can achieve noninvasive, spatiotemporally precise targeting of any brain region in small animals [7], large animals [2], and even humans [8]. Sonogenetics was first demonstrated in 2015 using *C. elegans*, in which mechanosensitive TRP-4 ion channel expression in neurons in combination with microbubbles evoked behavioral changes upon ultrasound stimulation [9]. Since the first demonstration of sonogenetics, many other mechanosensitive ion channels and proteins have been proposed to sensitize cells to ultrasound stimulation *in vitro*, including TREK1, TREK2, TRAAK [10], MscL [11], Piezo1 [12], MEC-4 [13], prestin [14], TRPA1 [15], TRPC1, TRPP2, and TRPM4 [16]. Recently, multiple studies demonstrated the feasibility of sonogenetics to modulate mouse behavior *in vivo* using ultrasound-sensitive probes such as prestin [17], MscL G22S [18], and TRPA1 [19].

Ultrasound propagation in tissue can generate not only mechanical effects but also thermal effects. The transient receptor potential vanilloid 1 (TRPV1) ion channel is extremely sensitive to temperature and has a thermal activation threshold of approximately 42°C, which is only a few degrees above the physiological body temperature of many mammals [20]. Such an activation temperature allows TRPV1 to be closed at the physiological body temperature and open upon sufficient heating to ~42°C. Because of these unique features, TRPV1 has been used to develop genetics-based neuromodulation techniques, such as magneto-thermogenetics [21,22] and photothermal genetics [23]. TRPV1-mediated sonogenetics was recently developed to achieve noninvasive, cell-type specific neuromodulation. Our previous study demonstrated that TRPV1 is an ultrasound-sensitive actuator, and that TRPV1-mediated sonogenetics can control the motor behavior of freely moving mice by targeting a deep brain region, the striatum [24]. However, the capability of TRPV1-mediated sonogenetics in controlling mouse behavior by targeting the superficial brain area has not been demonstrated. Targeting superficial brain regions is challenging for TRPV1-mediated sonogenetics because the high absorption of ultrasound in the skull could increase the risk of overheating the cortex area directly underneath the skull [25,26]. This could increase the risk of undesirable neuromodulatory effects associated with heating and potential tissue damage when the temperature is high. Therefore, the objective of the current study was to assess the capability of TRPV1-mediated sonogenetics in evoking mouse motor behavior by targeting a superficial brain target – the motor cortex.

## Materials and methods

### Stereotaxic injection of virus

All animal procedures were performed under a protocol approved by the Washington University in St. Louis Institutional Animal Care and Use Committee (IACUC). C57/BL6 mice (female, 6-8 weeks old) were purchased from Charles River and housed in an animal facility under a 12 hour light-dark cycle. Adeno-associated viruses (AAV) were introduced to CaMKII-expressing neurons of the M2 cortex to overexpress TRPV1 ion channel. All surgeries were conducted under aseptic conditions. Mice were anesthetized with 2% isoflurane in oxygen at a rate of 1.0 L/min in an anesthetic chamber for induction and 1.5% isoflurane for maintaining anesthesia. Anesthetized mice were then fixed onto a stereotaxic frame (Kopf Instruments) using a bite bar and ear bars. Buprenorphine SR (1.0 mg/kg) was administered subcutaneously for pre-operative and post-operative pain management. The head was shaved and was rubbed with skin disinfectant (Hibiclens). An incision was made on the scalp, the skin was retracted, and the periosteum was removed. A small hole was drilled through the skull (−1.0 mm ML, +2.5 mm AP, −1.0 mm DV), and a micro-injector (Nanoject II, Drummond Scientific) was inserted into the motor cortex. 1200 nL of TRPV1 virus (1.4e12 vg/mL) was introduced at a rate of 64 nL/min. 1000 nL of control virus (3.2e12 vg/mL) was introduced to approximately match the viral genome copy numbers delivered to the motor cortex. After injection, the micro-injector was slowly removed, the hole was filled with bone wax, and the scalp was sutured. Mice were housed for at least 4 weeks to facilitate sufficient virus expression before further treatments were conducted.

### Wearable FUS device

A wearable FUS device was used to stimulate the motor cortex of freely moving mice, and the design is described in the first report of TRPV1-mediated sonogenetics [24]. In brief, the wearable FUS device consisted of two parts: a FUS transducer and a base plate. The FUS transducer was made of a lead zirconate titanate (PZT) ceramic resonator (DL-43, DeL Piezo Specialties) encapsulated by a 3D-printed housing. The PZT ceramic resonated at a frequency of 1.5 MHz and had an aperture of 10 mm and a radius of curvature of 10 mm. The wearable FUS transducer was plugged into the base plate, a 3D printed circular adapter attached to the mouse skull. When the FUS transducer was plugged into the base plate, the wearable FUS device was stabilized on the mouse head.

Each component of the wearable FUS device was specifically designed to target the motor cortex. The base plate was designed with a hole in its geometric center to facilitate the alignment of base plate to the medial-lateral and anterior-posterior coordinates of the motor cortex. The height of the FUS transducer housing was designed to align the FUS focus to the dorsal-ventral coordinates of the motor cortex. The entire wearable FUS device was calibrated by a hydrophone (HGL-200, Onda). The full width half-maximum of the FUS focal region was approximately 0.9 mm and 2.5 mm in the lateral and axial directions, respectively.

### Attachment of FUS transducer base plate to the mouse skull

Four to five weeks after virus injection, mice were again anesthetized with isoflurane (2% for induction, 1.5% for maintenance), fixed in a stereotaxic frame, and subcutaneously administered with Buprenorphine SR (1.0 mg/kg). A piece of the scalp was removed, the periosteum was removed, and the drilled hole from the intracranial injection of AAV was identified and accentuated with a marker. The custom designed base plate was 3D-printed and glued onto the skull using dental adhesives (Metabond) with the center of the base plate aligned to the pre-drilled hole. The mice were housed for a week to facilitate sufficient recovery before performing behavior experiments.

### FUS stimulation with behavior recording

Prior to the behavior test, mice were adapted to the behavior recording environment by placing the mouse in the behavior testing arena with the power amplifier turned on. During the behavior recording, mice were lightly anesthetized with isoflurane (1% induction and maintenance). The base plate on the mouse and the wearable ultrasound transducer were both sufficiently filled with degassed ultrasound gel (Aquasonics). The wearable transducer was then securely plugged into the base plate of the mouse, and the mouse was then placed in a circular arena on a heating pad for 30 min to allow the mouse body temperature to recover from any possible anesthesia effects. The heating pad was then removed, and the mouse was allowed to habituate for 15 min in the actual behavior test arena.

During the recording period, focused ultrasound was applied at a frequency of 1.5 MHz, duty cycle of 40%, PRF of 10 Hz, and 15 s total sonication duration with 185 s inter-stimulation interval for a total of 5 stimulations. The onset and offset of the ultrasound pulse was smoothed to avoid possible auditory effects [27]. The acoustic pressures used in the study were 0, 0.7, and 1.1 MPa to investigate the effect of pressure on locomotor behavior outcomes. Custom MATLAB software was used to control when ultrasound was applied via an Arduino Uno. A red LED attached to the Arduino Uno would turn on when ultrasound was applied to precisely synchronize mouse behavior to each focused ultrasound stimulation. In each group, mice were given five consecutive focused ultrasound stimulations at one pressure.

### Behavioral analysis

Mice were recorded using a camera (Logitech C920X, 30 fps) before, during, and after each focused ultrasound stimulation. During the recording session, each video is simultaneously processed using Bonsai to quantify the positional coordinates and the angular orientation of the mice. After conducting recordings, data were processed using a custom MATLAB script to compute the average angular velocities upon FUS stimulation at different acoustic pressures.

### Immunohistological analysis

Approximately 90 minutes after the last FUS stimulation, TRPV1- and TRPV1+ mice from each acoustic pressure stimulation group were sacrificed via transcardial perfusion with 1x PBS solution for the evaluation of TRPV1 expression, c-Fos expression, and safety of sonogenetics via inflammatory and apoptotic markers (GFAP, Iba1, and Caspase-3). A sacrifice time of 90 minutes post stimulation was chosen to visualize the peak expression of c-Fos [28]. This time was also suitable to visualize any rapid recruitment of inflammatory and apoptotic markers at the FUS stimulation site [29]. The brains were fixed in 4% w/v paraformaldehyde in 1x PBS solution overnight and were transferred to 15% and 30% w/v sucrose in 1x PBS for the following two days, respectively. The brain tissue was embedded in a cryomold with Optimal Cutting Temperature medium (Scigen) to generate 10 μm thick coronal brain slices affixed on a glass slide.

For evaluation of TRPV1 and c-Fos expression, slides with brain tissue were stained with anti-TRPV1 antibody (Novus Biologicals, 1:200), anti-c-Fos antibody (Cell Signaling, 1:1000), and Nissl stain (Invitrogen, 1:100). TRPV1 and c-Fos were visualized using Alexa Fluor 594 and 488 secondary antibody (Jackson ImmunoResearch, 1:400), respectively. For cellular safety evaluation, slides with brain tissue were stained with anti-GFAP antibody (Abcam, 1:1000), anti-Iba1 antibody (Wako, 1:1000), or anti-Caspase-3 antibody (Cell Signaling, 1:2500), as well as DAPI mounting medium (Vector). GFAP, Iba1, and Caspase-3 cells were visualized using Alexa Fluor 488 secondary antibody (Jackson ImmunoResearch, 1:400). Cell counts were computed in the motor cortex using QuPath (University of Edinburgh). The viral spread was quantified by drawing a region that encapsulated all the TRPV1+ neurons. TRPV1+ and c-Fos+ neuron cell densities were calculated by counting the total number of positively stained Nissl cells over the motor cortex region. GFAP, Iba1, and Caspase-3 cell counts were calculated by counting the total number of positively-stained DAPI cells over the motor cortex region.

### Statistics

Statistical tests were conducted using GraphPad. Data were analyzed using either a two-tailed t-test or repeated measures ANOVA with either Bonferroni’s post-hoc test (to compare row- and column-wise groups) or Dunnett’s post-hoc test (to compare to a control group). Statistical differences were considered significant whenever *p*□<□0.05. All graphs presented results as the mean□±□standard error of the mean (SEM).

## Results

We intracranially injected adeno-associated virus (AAV) to the mouse motor cortex (M2) to express TRPV1 primarily in excitatory neurons under the CaMKII promoter (**Fig. 1a**). These mice are referred to as TRPV1+ mice. Control mice were injected with TRPV1-virus, referred to as TRPV1-mice. After sufficient virus expression, a wearable FUS transducer was attached to the mouse head. Mouse locomotion was assessed before, during, and after FUS sonication in an open-field behavior test arena. FUS sonication was targeted at the motor cortex using the same coordinates as the virus injection by mechanically aligning the FUS device to the craniotomy from the virus injection. FUS was applied with a center frequency of 1.5 MHz, a pulse repetition frequency (PRF) of 10 Hz, a duty cycle (DC) of 40%, acoustic pressures of 0.7 and 1.1 MPa, and a burst duration (BD) of 15 s with an inter-stimulation interval (ISI) of 185 s for a total of five stimulations (**Fig. 1b**). Mice were sacrificed after the behavior test to evaluate the expression of TRPV1, the activation of neurons (c-Fos), and safety of sonogenetics.

**Fig. 1.**
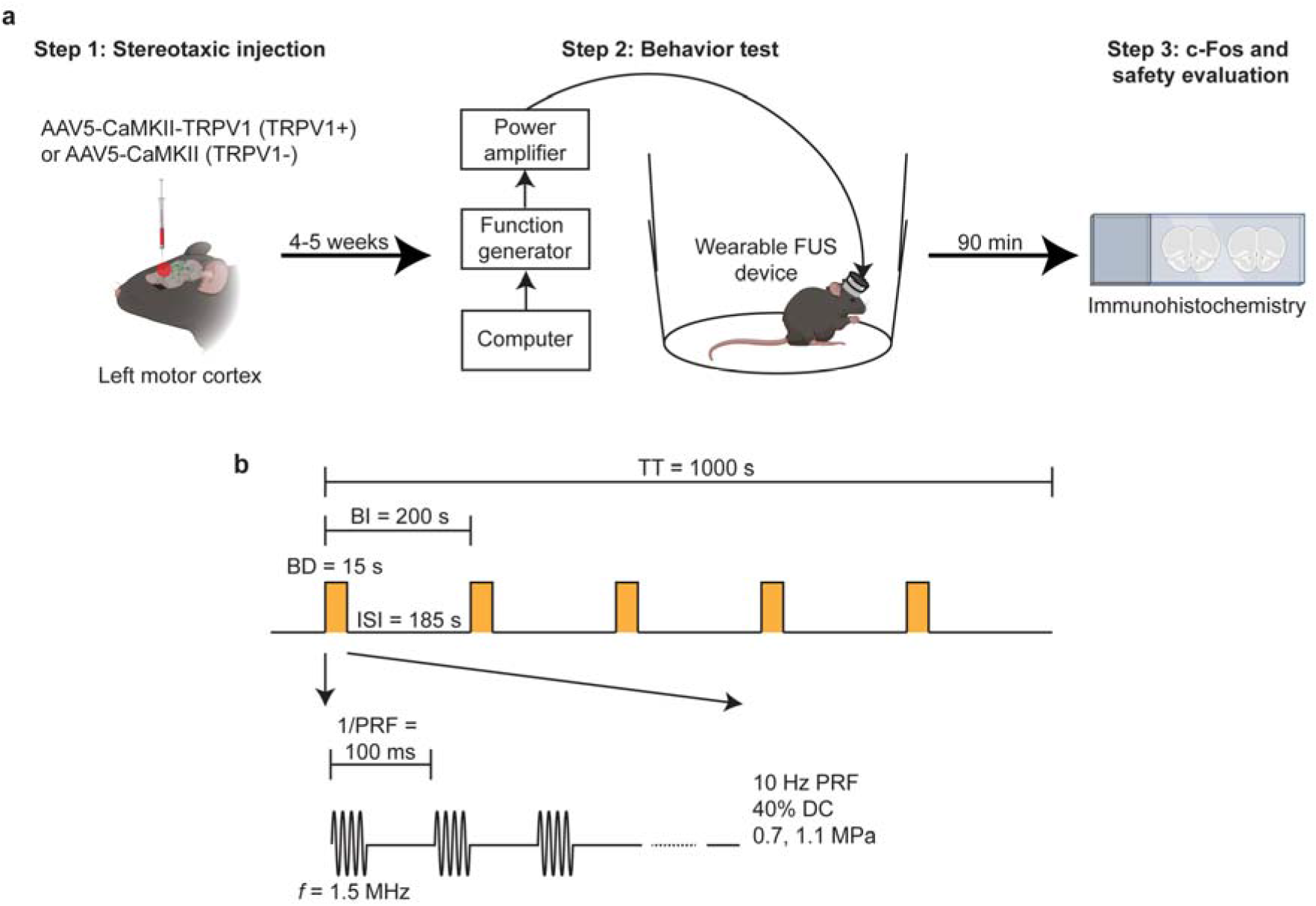
Experimental setup. (a) Experimental timeline. The study begins with intracranial injection of adeno-associated virus encoding TRPV1 (TRPV1+ mice). Control mice were injected with a control viral vector (TRPV1-mice). After 4-5 weeks, a wearable FUS device was installed onto the mouse skull to target the same location where the viral vectors were injected. The stimulation apparatus consists of a computer, function generator, and power amplifier to apply FUS to the wearable FUS device. Approximately 90 minutes after the final stimulation, mice brains were harvested for c-Fos and safety analyses. (b) Schematic of the ultrasound waveform used during the behavior test. FUS was applied with a center frequency of 1.5 MHz, a pulse repetition frequency (PRF) of 10 Hz, and a duty cycle (DC) of 40%. The burst duration (BD) was 15 s with an inter-stimulus interval (ISI) of 185 s, for a total of five stimulations with a total time (TT) of 1000 s.

### Characterization of exogenous TRPV1 expression in the motor cortex

We first describe the virus expression level of TRPV1 in the motor cortex. The brains of TRPV1+ mice were harvested, sectioned, and co-stained with anti-TRPV1 antibody and Nissl to evaluate the expression profile of TRPV1 in cortical neurons of the motor cortex. A representative brain slice of a TRPV1+ mouse illustrated that TRPV1 expression was primarily confined to the motor cortex (**Fig. 2a**). As expected, the contralateral non-injection site did not express any TRPV1 in cortical neurons of the motor cortex. The viral spread of TRPV1 in the cortex was 1.06 ± 0.07 mm^2^, and the density of neurons in the motor cortex that were virally transduced to express TRPV1 was 66.7 ± 4.0 cells/mm^2^ (**Fig. 2b-c**). The proportion of neurons in the motor cortex that expressed TRPV1 cells was 5.16 ± 0.43% (**Fig. 2d**). Higher magnification of the virus transduction region showed that TRPV1 expression was largely confined to neurons, in which 83.4 ± 3.0% of the cells transfected with TRPV1 were neurons (**Fig. 2e**). These data demonstrate the feasibility of exogenous TRPV1 expression in the motor cortex region and lay the foundation to facilitate TRPV1-mediated sonogenetic control of motor cortex behaviors.

**Fig. 2.**
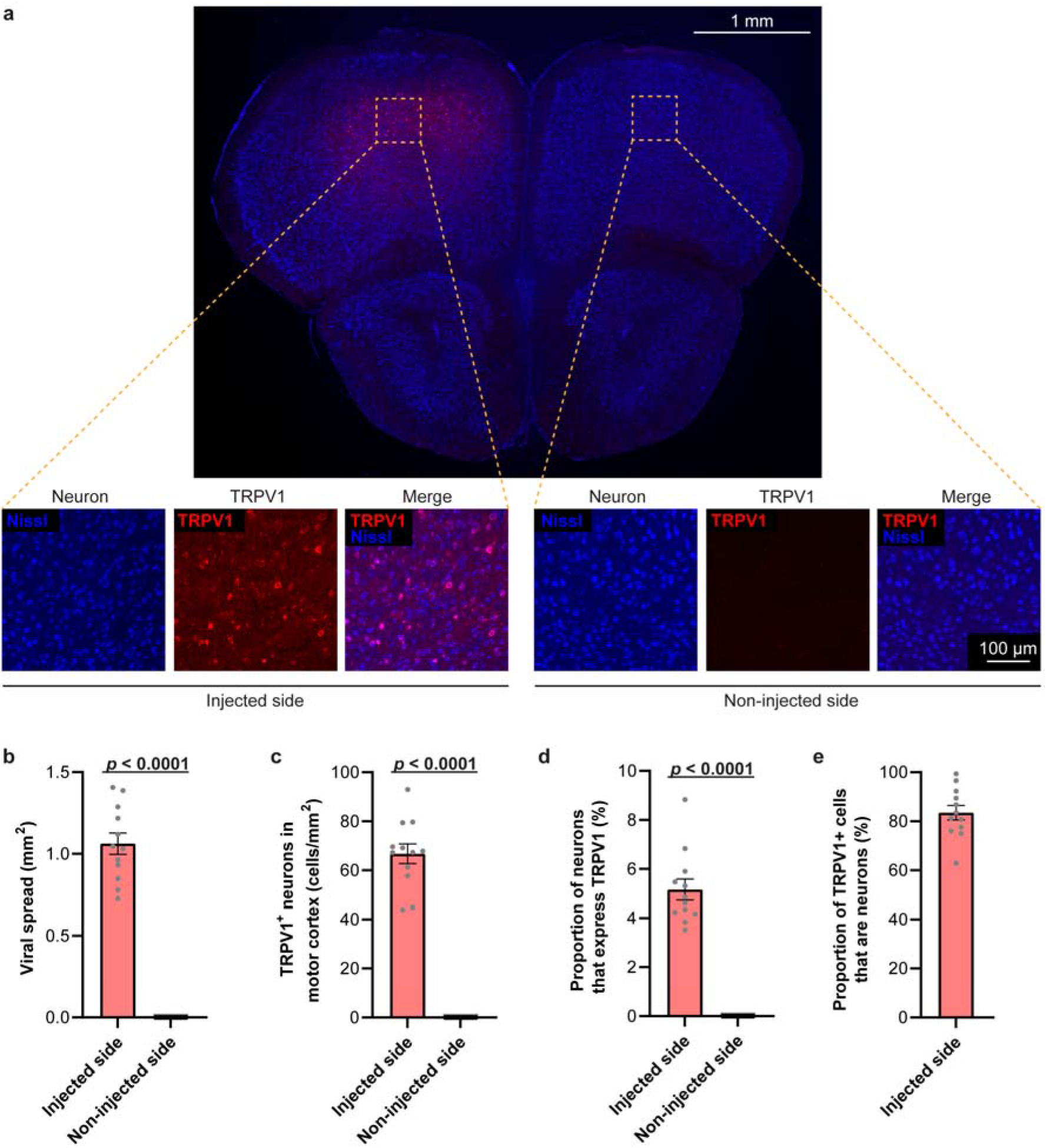
Characterization of exogenous TRPV1 expression in the motor cortex. (a) Representative immunofluorescence image of TRPV1+ mouse brain slice that has been stained with anti-TRPV1 antibody (red) and Nissl dye (blue) (Scale bar = 1 mm). The yellow box corresponds to a higher magnification image (Scale bar = 100 μm). Quantification of (b) the viral spread of TRPV1 expression, (c) the density of TRPV1+ neurons in the motor cortex, (d) the proportion of TRPV1+ neurons in the motor cortex, and (e) the proportion of TRPV1+ cells that are neurons. Data are reported as mean ± SEM.

### TRPV1-mediated sonogenetic neuromodulation within the motor cortex alters locomotor behavior

We recorded the locomotor behavior of TRPV1- and TRPV1+ mice with the application of FUS at the motor cortex to assess the ability of TRPV1-mediated sonogenetics in modulating locomotor behavior. Representative locomotor behavior of TRPV1- and TRPV1+ mice with and without FUS are shown as position traces (**Fig. 3a; TRPV1-, Movie S1; TRPV1+, Movie S2**). FUS sonication at 0.7 MPa did not evoke considerable motion in TRPV1-mice compared to that before FUS. In TRPV1+ mice, however, FUS stimulation did evoke rotational behavior around the behavior testing arena, which was not observed before FUS. During the FUS sonication periods (shown by the highlighted yellow bars), TRPV1+ mice displayed rotational behavior indicated by changes in angular displacement and angular velocity (**Fig. 3, 3c**). In contrast, TRPV1-mice did not demonstrate any rotational bias upon FUS sonication.

**Fig. 3.**
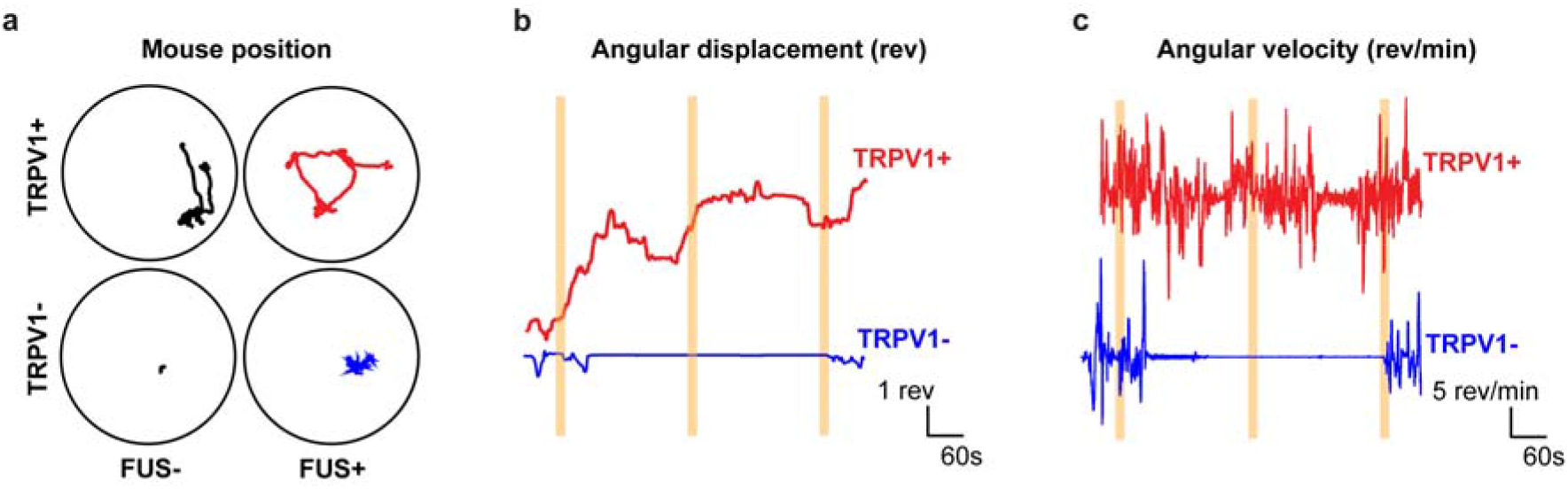
Sonogenetics with TRPV1 evokes rotational behavior at 0.7 MPa. (a) Representative position plots of TRPV1+ and TRPV1-mice with and without the application of one FUS stimulation. Representative plots of (b) the angular displacement over time and (c) angular velocity over time. The yellow bars correspond to the application of FUS at an acoustic pressure of 0.7 MPa.

We then compared the average angular velocities of TRPV1- and TRPV1+ mice with FUS at acoustic pressures of 0.7 and 1.1 MPa. In the TRPV1+ mice group, FUS stimulation at 0.7 MPa evoked a significant increase in angular velocity (0.86 ± 0.23 rev/min) compared to the sham stimulation at 0 MPa (−0.22 ± 0.25 rev/min), indicating that TRPV1+ mice displayed a preference to rotate in the direction contralateral to the stimulation site (**Fig. 4**; ~4-fold increase, *p* = 0.026, two-way repeated measures ANOVA with Bonferroni’s post-hoc test). In contrast, FUS stimulation at 0.7 MPa did not evoke any significant angular velocity changes in the TRPV1-mice group relative to the sham stimulation (0.7 MPa: −0.20 ± 0.31 rev/min; 0 MPa: −0.17 ± 0.21 rev/min). These findings indicate that TRPV1-mediated sonogenetics at 0.7 MPa can achieve circuit-specific control of locomotor behaviors in the motor cortex. Increasing the acoustic pressure to 1.1 MPa did not evoke significant changes in angular velocity compared to the sham stimulation at 0 MPa in TRPV1+ mice (1.1 MPa: 0.36 ± 0.53 rev/min). However, sonogenetics stimulation at 1.1 MPa of TRPV1+ mice achieved a significantly higher angular velocity than that obtained by FUS sonication at 1.1 MPa of TRPV1-mice (*p* = 0.036). Although not statistically significant, FUS sonication at 1.1 MPa of TRPV1-mice evoked an increase in the average angular velocity in the ipsilateral direction (−0.88 ± 0.36 rev/min) compared with those at 0 MPa and 0.7 MPa. These findings suggest that FUS stimulation at 1.1 MPa alone (without TRPV1) potentially induced neuromodulation effects and generated a confounding impact on TRPV1-mediated sonogenetics at this high-pressure level.

**Fig. 4.**
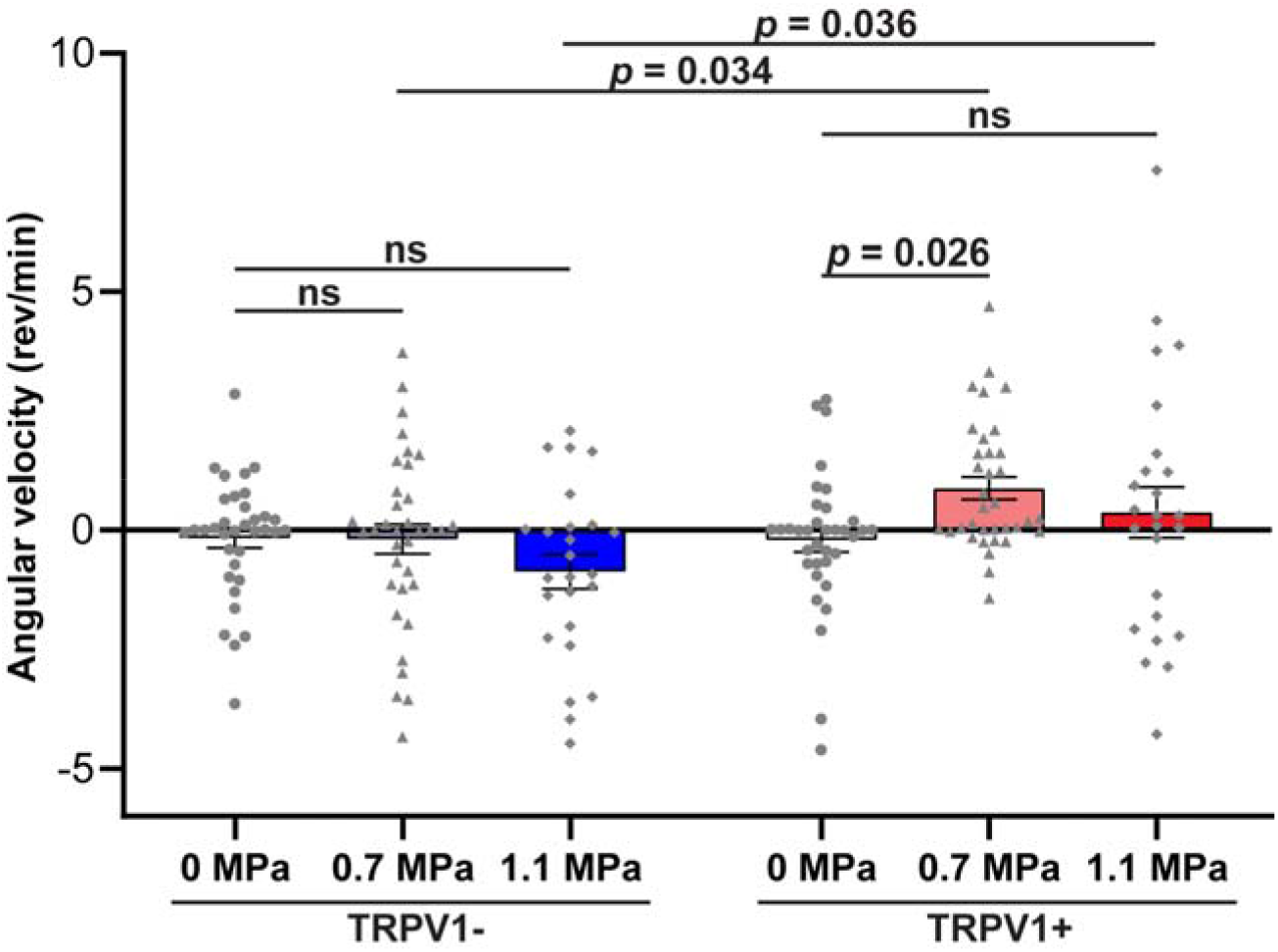
TRPV1-mediated sonogenetics at 0.7 MPa facilitates direction-specific locomotor control. Summary plot of the average angular velocity for TRPV1- and TRPV1+ mice at 0, 0.7, and 1.1 MPa FUS sonications. Angular velocity values greater than zero correspond to contralateral rotations (clockwise), while angular velocity values less than zero correspond to ipsilateral rotations (counter-clockwise). Each point represents one stimulation. Data are reported as mean ± SEM. Statistical analysis was conducted using two-way repeated measures ANOVA with Bonferroni post-hoc test.

### TRPV1-mediated sonogenetics activates cortical neurons on the cellular level

To provide a secondary readout for successful modulation of the motor cortex using sonogenetics, we sacrificed the mice 90 minutes after the final stimulation and used immunohistochemical staining to analyze c-Fos expression levels of TRPV1+ and TRPV1-mice. Representative fluorescent images of TRPV1- and TRPV1+ brains stimulated at different acoustic pressures demonstrate that sonication at both 0.7 MPa and 1.1 MPa elicited greater c-Fos expression levels in the motor cortex of TRPV1+ mice (**Fig. 5a**). Group analysis found that TRPV1+ mice showed enhancement in the number of c-Fos cells at both 0.7 MPa (231.5 ± 58.3 cells/mm^2^) and 1.1 MPa (332.1 ± 74.2 cells/mm^2^) compared to the unstimulated side (89.4 ± 22.2 cells/mm^2^), indicating activation of neurons in the motor cortex (**Fig. 5b**; 0.7 MPa: ~2.6-fold change, *p* = 0.011; 1.1 MPa: ~3.7-fold change, *p* < 0.0001; two-way ANOVA with Bonferroni’s post-hoc test). On the other hand, FUS stimulation at 0.7 MPa and 1.1 MPa did not evoke significant enhancements in c-Fos expression in TRPV1-mice (0 MPa: 70.1 ± 19.1 cells/mm^2^; 0.7 MPa: 51.8 ± 21.8 cells/mm^2^; 1.1 MPa: 144.5 ± 50.4 cells/mm^2^). While there is a slight potential increase in c-Fos expression in TRPV1-mice from FUS alone at 1.1 MPa compared to the unstimulated control, this relationship was not statistically significant (*p* = 0.34). These data demonstrate the ability of TRPV1-mediated sonogenetics to activate motor cortex neurons at the cellular level at both 0.7 MPa and 1.1 MPa.

**Fig. 5.**
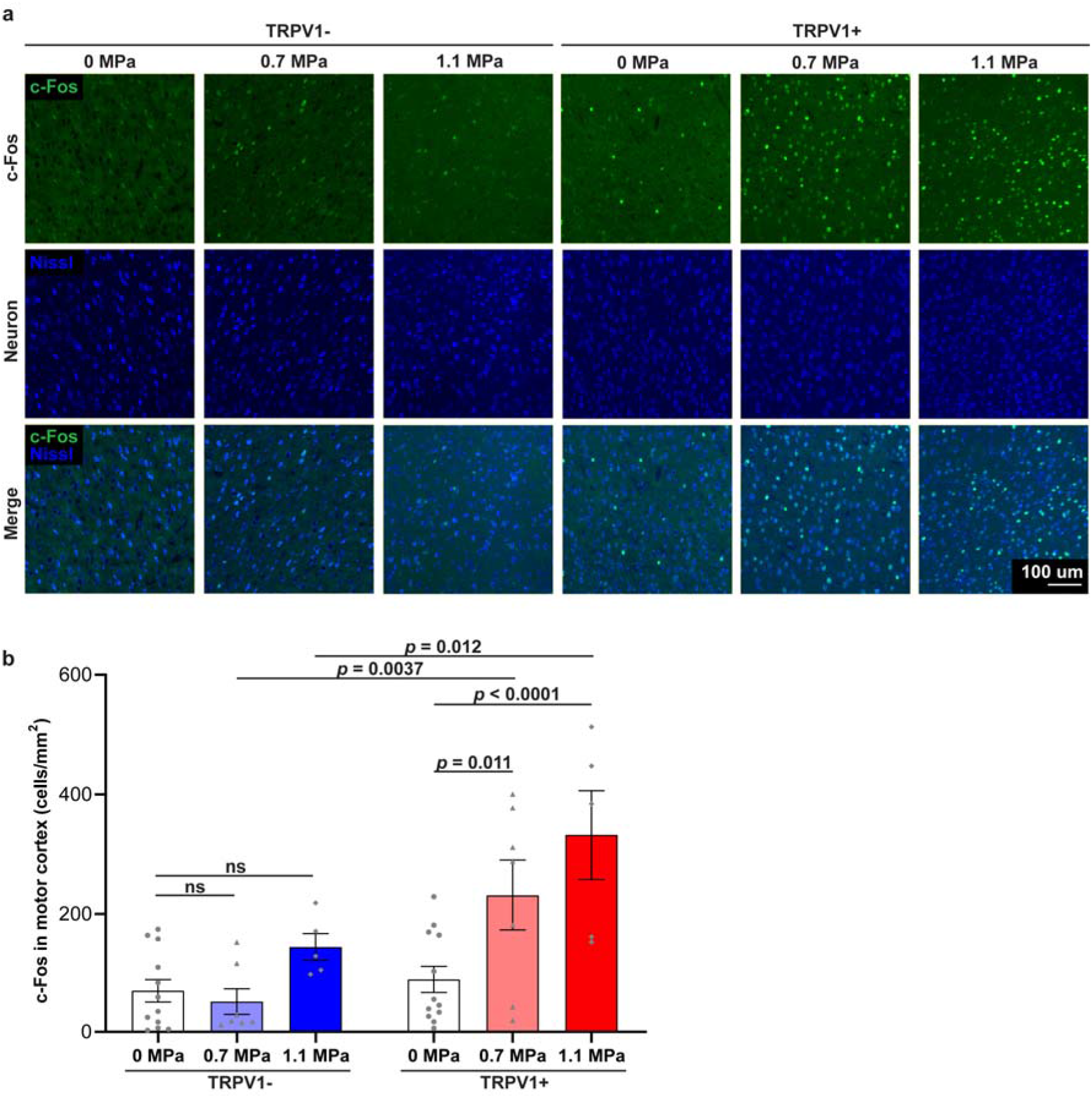
TRPV1-mediated sonogenetics activates cortical neurons at the cellular level. (a) Representative immunofluorescence images of TRPV1- and TRPV1+ mice brains stained with anti-c-Fos antibody (green) and Nissl dye (blue) at the unstimulated side, 0.7, and 1.1 MPa (Scale bar = 100 μm). (b) Quantification of the c-Fos+ neuron count in the motor cortex. FUS sonication at 0.7 and 1.1 MPa enhanced the number of c-Fos expressing neurons in the motor cortex in TRPV1+ mice. Data are reported as mean ± SEM. Statistical analysis was conducted with two-way repeated-measures ANOVA with Bonferroni’s posthoc test.

### Inflammatory and apoptotic responses in the brain are not engaged by TRPV1-mediated sonogenetics

Gross pathology of the mice skull and brain stimulated at 0.7 MPa showed no signs of damage (Fig. 6a). In contrast, bleeding was consistently observed in the meninges between the skull and the brain at 1.1 MPa. Furthermore, we used Nissl to stain for signs of neuronal damage, GFAP and Iba1 to stain for signs of inflammation, and Caspase-3 to stain for signs of apoptosis (Fig. 6b). Using the non-injection and non-stimulated side of both TRPV1+ and TRPV1-mice as the control, there were no significant differences in any of the protein marker expression levels in the mouse brain at 0.7 MPa or 1.1 MPa (Fig. 6c; one-way repeated measures ANOVA with Dunnett’s post-hoc test). Both gross pathology and immunohistological analysis of inflammatory and apoptotic markers showed that TRPV1-mediated sonogenetics at 0.7 MPa enables safe neuromodulation, while damage at the meninges was associated with sonogenetics at 1.1 MPa.

**Fig. 6.**
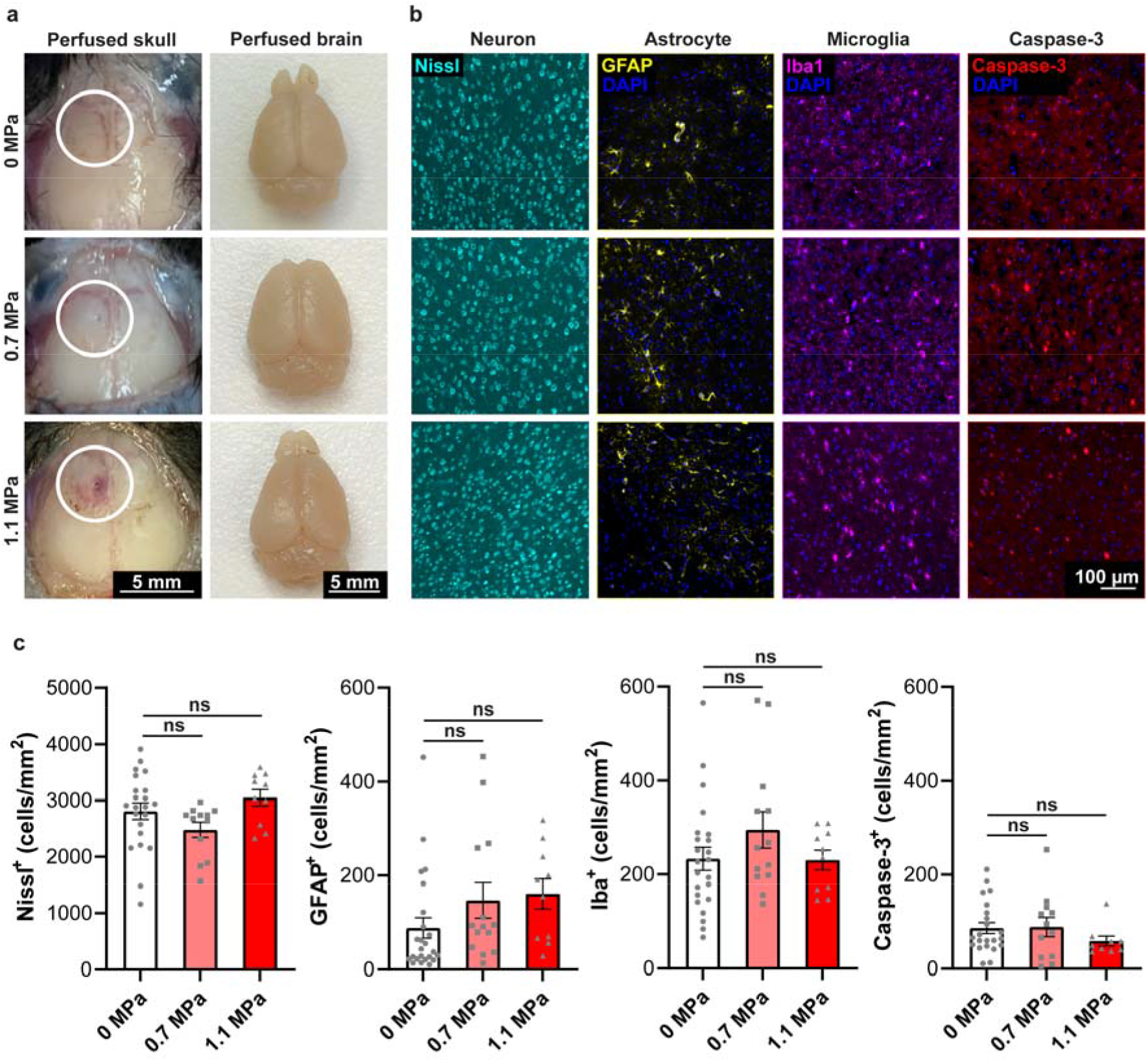
TRPV1-mediated sonogenetics at 0.7 MPa did not show signs of inflammation or apoptosis. (a) Representative gross pathology images of the mice skull and brain 90 minutes after the last FUS sonication at 0, 0.7, and 1.1 MPa. The first column of images shows the intact skull of the perfused mouse, and the second column of images shows the intact brain of the perfused mouse (scale bar = 5 mm). (b) Representative immunofluorescence images of mice brains stained with Nissl dye (cyan), anti-GFAP antibody (yellow), anti-Iba1 antibody (magenta), and anti-Caspase-3 antibody (red) at 0, 0.7, and 1.1 MPa (scale bar = 100 μm). (c) Summary of neuron, astrocyte, microglia, and caspase-3 cell counts in the mice motor cortex after FUS sonication at 0, 0.7, and 1.1 MPa. Data are reported as mean ± SEM. Statistical analysis was conducted with one-way repeated-measures ANOVA with Dunnett’s post-hoc test.

## Discussion

Sonogenetics is a rapidly emerging technique that enables noninvasive, cell-type specific neuromodulation with high spatiotemporal resolution. This study demonstrates the capability of TRPV1-mediated sonogenetics to modulate behavior in freely moving mice.

Previous studies have reported sonogenetic-enabled neuromodulation in mice by activating mechanosensitive ion channels and proteins, such as prestin [17], MscL G22S [18], and TRPA1 [19], using FUS-induced mechanical effects. They observed neuron activation based on c-Fos staining, as well as motor responses in head-fixed anesthetized mice based on electromyography, but did not report induction of real-time behavior modulation in freely moving mice. Different from these studies, the current study used thermosensitive ion channel TRPV1-mediated sonogenetics and achieved successful behavior modulation in freely moving mice. Our previous study demonstrated successful locomotor behavior modulation by TRPV1-mediated sonogenetics in freely moving mice by targeting a deep brain region, the striatum [24]. Here we demonstrated that this technique could modulate locomotor behaviors via a superficial brain target (motor cortex), expanding the application of TRPV1-mediated sonogenetics to include both superficial and deep brain targets.

Previous studies have also reported successful neuromodulation using TRPV1-mediated neuromodulation via combining TRPV1 with different external stimulation modalities. TRPV1-mediated magnetothermal-genetics targeting both superficial and deep brain targets were previously reported [21,22]; however, magnetic nanoparticles need to be injected into the brain to convert energy from an alternating magnetic field to heat for TRPV1 activation. Recently, TRPV1-mediated photothermal genetic stimulation was reported, which combines an injection of nanoparticles with near-infrared light to generate heat for TRPV1 activation [23]. TRPV1-mediated magnetothermal and photothermal genetic modulation of the motor cortex induced increases in the angular speed of freely moving mice in the contralateral direction, which was consistent with the results of this study at 0.7 MPa sonication. However, both existing techniques require an additional component of “energy-converting” nanoparticles that were directly injected into brain tissue. The injection process poses inflammation and ischemia risks, and the presence of nanoparticles in the brain possesses immunogenic and biocompatibility concerns. Since FUS-mediated heating does not require the injection of nanoparticles, it provides a powerful alternative approach to achieving TRPV1-mediated genetic neuromodulation.

We show that TRPV1-mediated sonogenetics successfully evoked motor behavior by targeting the superficial brain target with carefully selected ultrasound parameters. Ultrasound parameters must be selected to achieve successful behavior control without causing any detectable tissue damage. Based on behavior, c-Fos, and safety analyses, TRPV1-mediated sonogenetics at 0.7 MPa met this requirement. However, increasing the pressure to 1.1 MPa did not evoke statistically significant changes in angular velocity in TRPV1+ mice relative to the sham sonication at 0 MPa, but evoked significant changes compared to FUS sonication of TRPV1-mice at 1.1 MPa. FUS sonication at 1.1 MPa in TRPV1-mice showed a trend, although not significant, to evoke ipsilateral rotations. These findings suggested that FUS stimulation at 1.1 MPa alone could impact animal behavior. It was interesting to find that damage to the meninges was observed at 1.1 MPa, although no damage to the brain tissue was clearly detected. Damage to the meninges was due to its proximity to the skull. The high skull absorption of ultrasound at 1.1 MPa caused thermal-induced damage to the meninges. Therefore, ultrasound parameter selection must be carefully selected when performing TRPV1-mediated sonogenetics to achieve effective and safe neuromodulation.

## Conclusion

In conclusion, our findings demonstrated the feasibility and safety of using TRPV1-mediated sonogenetics to modulate locomotor behaviors by targeting the motor cortex. Combined with our previous report on TRPV1-mediated sonogenetics for behavior modulation by targeting the deep brain region, our present study indicates that this technique can facilitate neuromodulation at the whole depth of the mouse brain.

## Acknowledgments

This work was supported by the NIH R01EB027223, R01EB030102, and R01MH116981.This work was also supported by the Hope Center Viral Vectors Core, the Alafi Neuroimaging Laboratory, the Hope Center for Neurological Disorders, and the NIH Shared Instrumentation Grant (S10 RR0227552) to Washington University.

## Data statement

The data that from this study are available from the corresponding author upon reasonable request.

